# Impairments in proprioceptively-referenced limb and eye movements in chronic stroke

**DOI:** 10.1101/2024.05.09.593341

**Authors:** Duncan T. Tulimieri, Amelia Decarie, Tarkeshwar Singh, Jennifer A. Semrau

## Abstract

**Background:** Upper limb proprioceptive impairments are common after stroke and can affect daily function. Recent work has shown that stroke survivors have difficulty using visual information to improve proprioception. It is unclear how eye movements are impacted to guide action of the arm after stroke. Here, we aimed to understand how upper limb proprioceptive impairments impact eye movements in individuals with stroke.

**Methods:** Control (N=20) and stroke participants (N=20) performed a proprioceptive matching task with upper limb and eye movements. A KINARM exoskeleton with eye tracking was used to assess limb and eye kinematics. The upper limb was passively moved by the robot and participants matched the location with either an arm or eye movement. Accuracy was measured as the difference between passive robot movement location and the active limb matching (Hand-End Point Error) or the active eye movement matching (Eye-End Point Error).

**Results:** We found that individuals with stroke had significantly larger Hand and Eye-End Point Errors compared to controls. Further, we found that stroke participants had proprioceptive errors of the hand and eye were highly correlated [r=0.67, p=0.001], a relationship not observed for controls.

**Conclusions:** Eye movement accuracy declined as a function of proprioceptive impairment of the more-affected limb, which was used as a proprioceptive reference. The inability to use proprioceptive information of the arm to coordinate eye movements suggests that disordered proprioception impacts integration of sensory information across different modalities. These results have important implications for how vision is used to actively guide limb movement during rehabilitation.

## Introduction

Proprioception, our sense of limb position and motion^1^, is critical for the production of accurate and well-controlled movements of the upper limb^2,3^. Recent work has established that proprioceptive impairments are common and are present in 30% - 80% of stroke survivors^4–8^. Despite this, assessment and treatment of proprioception is limited in many clinical environments resulting in a significant evidence-practice gap^9,10^ in neurorehabilitation. This gap is likely due to poor measurement tools^11,12^, a lack of understanding concerning the contributions of proprioception to motor output after stroke^13^, and mixed efficacy for treatment of proprioception after stroke^14^.

While impaired proprioception can have considerable functional impacts for stroke survivors^15^, our understanding of how impairments in proprioception affect motor output is incomplete. Notably, the proprioceptive system does not operate in isolation and works closely with the visual system to guide movement, improve accuracy, and plan future movements^16–18^. Recently, we found that ∼20% of stroke survivors were unable to improve proprioceptively-based movement output when given visual feedback of the arm, a result also replicated in children with cerebral palsy ^19–22^. This is surprising, as using visual guidance of the limb is common practice during neurorehabiliation^23,24^. Further, this suggests that not only is proprioception impaired, but that the integration of multiple sensory systems may be compromised for some stroke survivors, resulting in a disconnect between the perceived position of the limb and visual guidance of the limb.

The current study aims to examine how proprioceptive impairments can potentially impact other sensory systems in stroke survivors, namely the visual system. Using a robotic proprioceptive matching task where the robot moves the more-affected arm which serves as a reference of limb position sense. Participants then mirror-matched the felt position of the more-affected limb with a movement of the less-affected arm or an eye movement. We hypothesized that proprioceptive impairments of the arm that occur as a result of stroke can negatively impact how proprioceptive information is used to generate eye movements. Here, we quantify proprioceptive accuracy of limb location as the comparison between end-point accuracy of the passively moved limb (more-affected) and the active movement location of the less-affected limb, and proprioceptive accuracy of eye movements as the difference in end-point accuracy of the passively moved limb (more-affected) and eye fixation location. We predicted that individuals with chronic stroke would have greater end-point error for limb-referenced eye movement locations compared to age-matched control participants.

## Methods

### Participants

A total of 40 participants (age-matched controls: N=20 and stroke participants: N=20) were included in this study. Participants were included if they were 18 years of age or older and had normal or corrected-to-normal vision. Participants were excluded if they had a history of disease that impacted sensation (e.g., peripheral neuropathy), prior history of upper body injury (e.g., rotator cuff tear), or history of neurological injury or disease (e.g., Parkinson’s Disease). Inclusion criteria specific to stroke participants were the occurrence of a single, unilateral stroke and currently being within the chronic (> 6 months) phase after stroke. To determine hand dominance, the participants completed the Edinburgh Handedness Inventory^25^. The following clinical measures were used to quantify and characterize several aspects of overall function in stroke participants, including upper limb motor function (Fugl-Meyer Upper Extremity Assessment^26^), functional ability (Functional Independence Measure^27^), proprioception (Thumb Localization Test)^28^, gross and fine motor control (Purdue Pegboard^29^), visuospatial neglect (Behavioral Inattention Test^30^), and cognitive function (Montreal Cognitive Assessment^31^). The study was approved by the University of Delaware Institutional Review Board and all participants provided written informed consent.

### Experimental Apparatus

The KINARM Exoskeleton Lab (BKIN Technologies, Kinston, ON, Canada) with an integrated EyeLink 1000 Remote Eye-Tracking System (SR Research EyeLink, Ottawa, ON, Canada) was used to collect kinematic data from arm and eye movements (Figure 1A)^32^. The participants’ arms were supported against gravity by the exoskeleton with their shoulders at ∼85° abduction. The exoskeleton allowed for two degrees of freedom in the horizontal plane: shoulder flexion/extension and elbow flexion/extension. After the arms of the exoskeleton were custom fit to each participant’s limb geometry, participants were wheeled into the horizontally mounted virtual reality display. Data from the robot was collected at 1000 Hz. The eye-tracker was mounted at the back of the virtual reality display, ∼80 cm in front of the participants eyes. The system tracked participant eye movements at 500 Hz and is capable of capturing eye movements as small as 0.25°. During the entirety of the experiment, vision of the arms and shoulders were occluded by a metal shutter and a bib.

**Figure 1:**
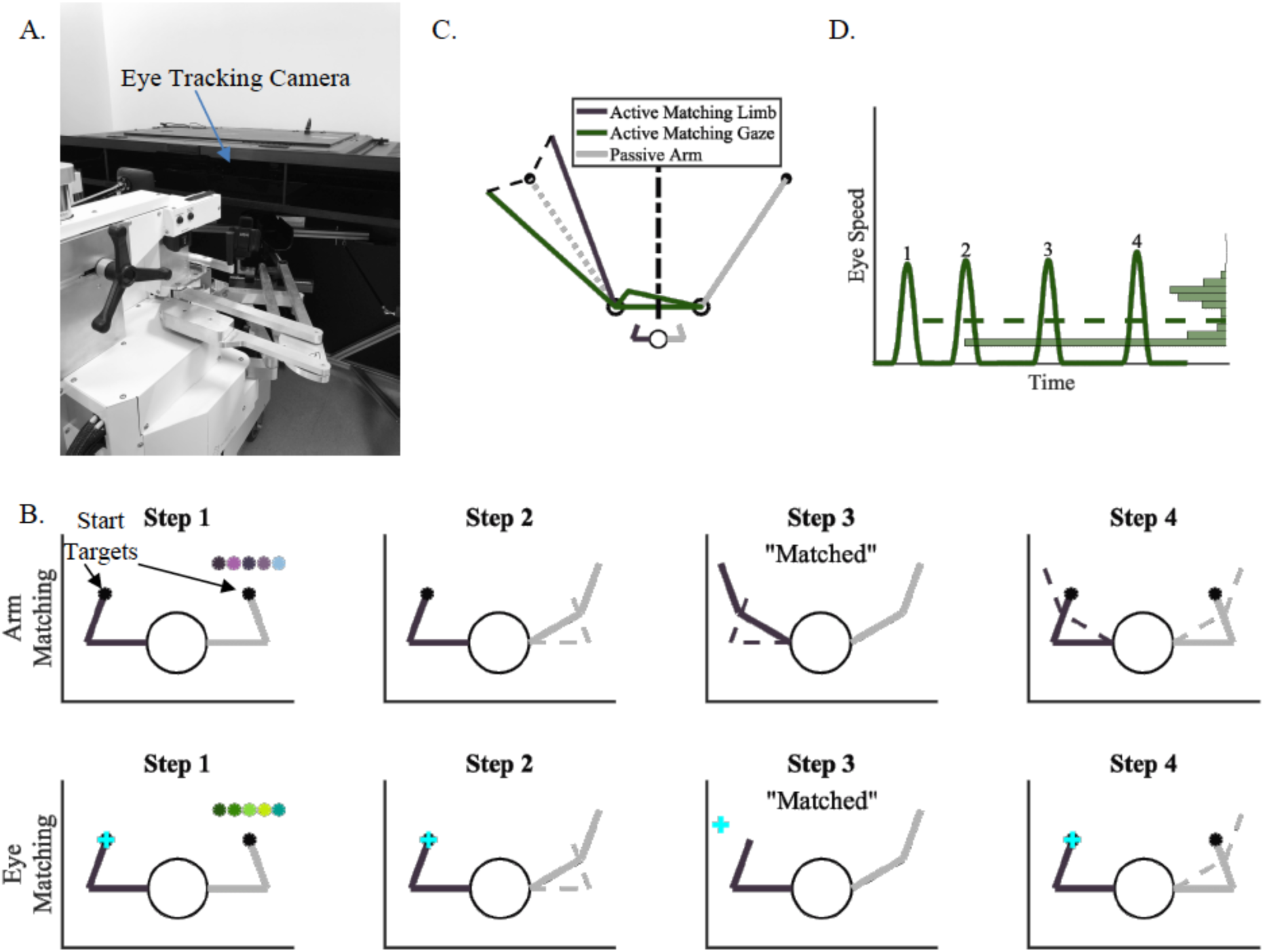
A). KINARM Robotic Exoskeleton. B). Single trial depiction for an arm matching trial (top row) and an eye matching trial (bottom row). Step 1: Participants started with both arms in the starting mirror-matched position. Step 2: The robot passively moved one arm (right arm) by the robot to one of five targets. Step 3: Participants mirror-matched the end position of the passive arm with an active arm movement (arm matching trials) or eye movement (eye matching trials) on the opposite side of the workspace. Step 4: The arms were returned to start position to begin the next trial. No visual information was available to participants in Steps 1-3. C). Modeled positional data from an arm and eye matching trial. End Point Error (black dashed lines) was calculated as the Euclidean distance from the final arm/eye position to the passive target position (black circle on the left). Black dashed-dotted line represents midline of workspace. D). Modeled velocity data from an arm and eye matching trial. Number of Saccades was calculated as the number of times eye movement velocity increased above and then decreased below their saccadic threshold as determined by individual velocity peak performance for eye tracking. Median Gaze Angular Velocity was calculated as the median (dashed line) of the distribution of eye movement velocities (histogram on right).

### Task Protocol

There were three types of trial blocks: arm-matching (30 trials/block), eye-matching (30 trials/block), and visual fixation (5 trials/block). The order of the main blocks (arm-matching and eye-matching) was counterbalanced within each group and repeated two times each. The supplemental block (visual fixation) was always presented between arm-matching and eye-matching blocks. All participants began the study with a familiarization block which consisted of 3 arm-matching trials, 3 eye-matching trials, and 5 visual fixation trials.

In the arm-matching blocks, participants viewed the fingertip location of both arms as a 1cm cursor to initially align both arms with the visual start target (2cm circle) on each side (Figure 1B, top row). At the start of the trial, all visual information was extinguished, and the robot passively moved one arm from the starting position to one of five unseen target locations. After the robot stopped moving, participants were instructed to make an arm movement to mirror-match the location of the passive arm using their opposite (active) arm. Participants verbally reported when they felt they were in a mirror-matched position and the operator ended the trial. The robot then moved both arms back to the starting position to begin the next trial. Before the start of the next trial, fingertip location was revealed for 500 ms to discourage proprioceptive drift and the position of both limbs was realigned with the start target^33^. Data from periods with visual feedback were not included in subsequent analyses. Participants performed 60 arm-matching trials in total, 30 trials per (2) arm-matching block.

The eye-matching blocks were similar to the hand-matching blocks as described above. Here, participants started with both arms in the start position and were instructed to use their eyes to fixate on the start position of the active arm (Figure 1B, bottom row). The robot then moved the passive arm from the start position to one of the five unseen end targets. After the passive arm stopped moving, participants were instructed to make an eye movement to mirror-match and fixate on the felt location of the passive arm. Participants verbally reported when they felt they were looking at the target and the trial was ended. Both arms were then returned to the starting position and the next trial began as described above for the hand-matching block trials. Participants performed 60 eye-matching trials in total, 30 trials per (2) eye-matching block.

Visual fixation blocks were used to ensure that the quality of eye tracking did not differ between the groups, as previous work has shown disordered eye movements in stroke survivors^34–36^. Participants were instructed to fixate their eye position on a series of visual stimuli that would appear on the screen one at a time (yellow 1 cm targets). One of five targets would appear on the screen and participants would maintain fixation for 1000 ms until the next target appeared. Participants made one fixation to each of the five targets for a total of five fixations during each block, for a total of 15 fixations across 3 blocks.

During the arm- and eye-matching blocks, one arm was passively moved by the robot. Within the control group, the passive arm was counterbalanced, and for stroke participants, the more-affected arm was always the arm that was passively moved by the robot.

### Data Analysis

#### Arm Movement Kinematics

Kinematic data from the upper limb was collected using the KINARM Robotic Exoskeleton. X and Y positional data was collected from both the active and passive arms and filtered using a double-pass third order filter with zero-lag. For analysis purposes, data from the active arm was mirrored across the x-axis to make direct comparisons to data from the passive, robotically-moved arm. The primary outcome measure of the hand-matching blocks was Hand-End Point Error (Hand-EPE). Hand-EPE was defined as the Euclidean distance from the position of the active hand at trial end to position of the passive hand at the same time (Figure 1C). A larger Hand-EPE indicated larger errors in estimating hand position of the passive arm.

#### Eye Movement Kinematics

Prior to analysis, eye movement data was cleaned, and saccadic thresholds were calculated (Supplemental Methods)^37^. Saccadic eye movements were classified as the times when the angular velocity of the eye increased above the participants’ saccadic threshold. Data were then analyzed to determine the position matching error of the eye during eye matching movements. Eye-End Point Error (Eye-EPE) was computed as the Euclidean distance from the position of the eye at the end of trial to the end point of the passive robotic movement at the same time (Figure 1C). A larger Eye-EPE indicated that the eye position was further from the ideal end position. Additional eye metrics were computed, including: Number of Saccades - the total number of times eye movement velocity increased above and decreased below their saccadic threshold during a trial (Figure 1D), Number of Workspace Crossings was defined as the number of times the eye position crossed the midline of the workspace (Figure 1C), and Median Angular Velocity was defined as the median of the distribution of the eye angular velocity and was used to characterize the speed of generated eye movements (Figure 1D).

### Statistical Analyses

To quantify differences between the groups for the measures described above, permutation tests and Common Language Effect Size (CLES)^38^ were used. First, to assess if the control group was older than the individuals post-stroke, we compared the ages of the control and stroke groups using a directional permutation test such that *H*_0_: *individuals post stroke* < *older controls*. To address our predictions regarding arm and eye behavior, directional (*H*_0_: *individuals post stroke* < *older controls*) tests were used to compare both primary outcomes (i.e., Hand-EPE and Eye-EPE) and non-directional (*H*_0_: *older controls* = *younger controls*) tests were used to compare all secondary outcomes (i.e., Number of Saccades, Number of Workspace Crossings, and Median Angular Velocity). Additionally, to explore the relationship between our primary outcome measures (Hand-EPE and Eye-EPE), Spearman’s correlations were bootstrapped 1,000,000 times to obtain a distribution of the correlation coefficients and p-values for each group.

## Results

We assessed position sense using the arm and eye as the end effectors in age-matched controls and stroke participants using a position-matching paradigm. No differences were observed between groups for age (Controls: 62.9 ± 11.1; p=0.25, Stroke: 66.65 ± 8.73 years, CLES=54.62, Table 1).

**Table 1:** Participant demographics. Summary statistics are presented either as mean ± standard deviation, or median [minimum, maximum].

|  | Stroke participants (N=20) |
| --- | --- |
| Age | $66.65 \pm 8.73$ |
| Sex | 8 M, 12 F |
| Dominant hand | 16 R, 4 L, 0 A |
| Months post-stroke | 60 [11, 199] |
| Side of body affected | 8 R, 12 L |
| FM-UE (66) | 62 [11, 66] |
| FIM (126) | 124 [97, 126] |
| TLT {0, 1, 2, 3} | 16, 2, 2, 0 |
| PPB | 4.50 [0, 10] |
| BIT-C (146) | $141.40 \pm 4.21$ |
| Field-cut | N = 6 |
| MoCA (30) | $26.05 \pm 2.61$ |

In general, stroke participants demonstrated qualitatively larger end point errors for both the arm-matching and eye-matching blocks compared to controls, as demonstrated by the single trial examples in Figure 2 (Figure 2 A-B, E-F). On average, the exemplar control participant had, on average, 3.3 ± 1.4 cm Hand-EPE and 7.2 ± 2.8 cm Eye-EPE. Stroke participant 1 (Figure 2 C, D) demonstrated performance closer to the control participant and had, on average, 5.3 ± 2.5 cm Hand-EPE and 9.2 ± 7.0 cm Eye-EPE. While stroke participant 2 (Figure 2 E, F) had, on average, 7.3 ± 3.0 cm Hand-EPE and 13.5 ± 7.4 cm Eye-EPE. When we examined average behavior for Hand-EPE and Eye-EPE between groups, we found that on average within each group, stroke participants had greater error and greater variability than control participants for both hand- and eye-matching blocks that was consistent across all targets (Controls: Hand-EPE: 3.3 ± 1.4 cm, Eye-EPE: 7.2 ± 2.8 cm; Stroke: Hand-EPE: 7.0 ± 4.1 cm, Eye-EPE: 11.0 ± 4.0 cm; Figure 3).

**Figure 2:**
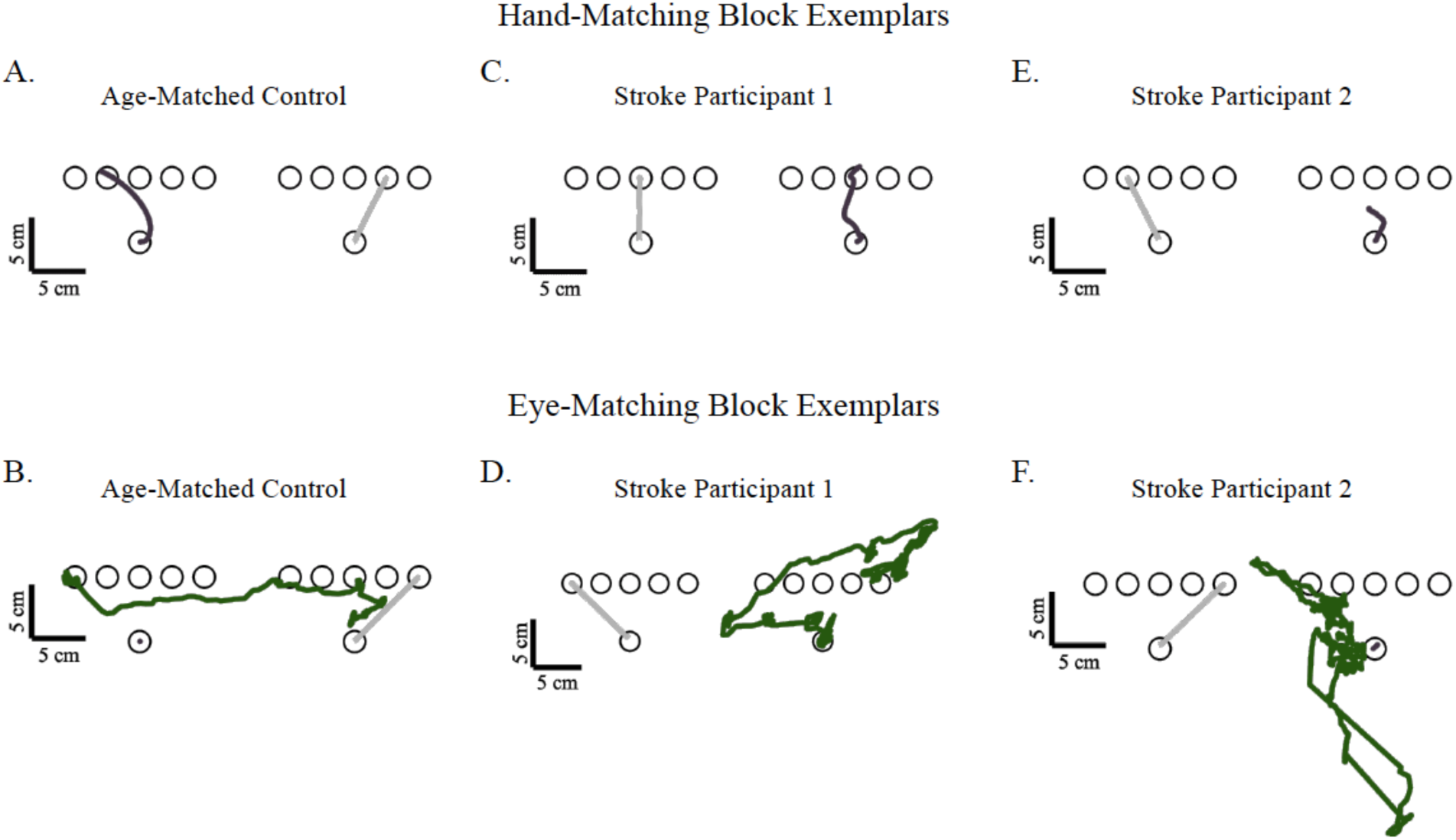
Exemplar data for a single trial for a control participant (A, B), stroke survivor with intact proprioception (C, D), and stroke survivor with impaired proprioception (E, F) for an arm matching (top row) and eye matching trial (bottom row).

**Figure 3:**
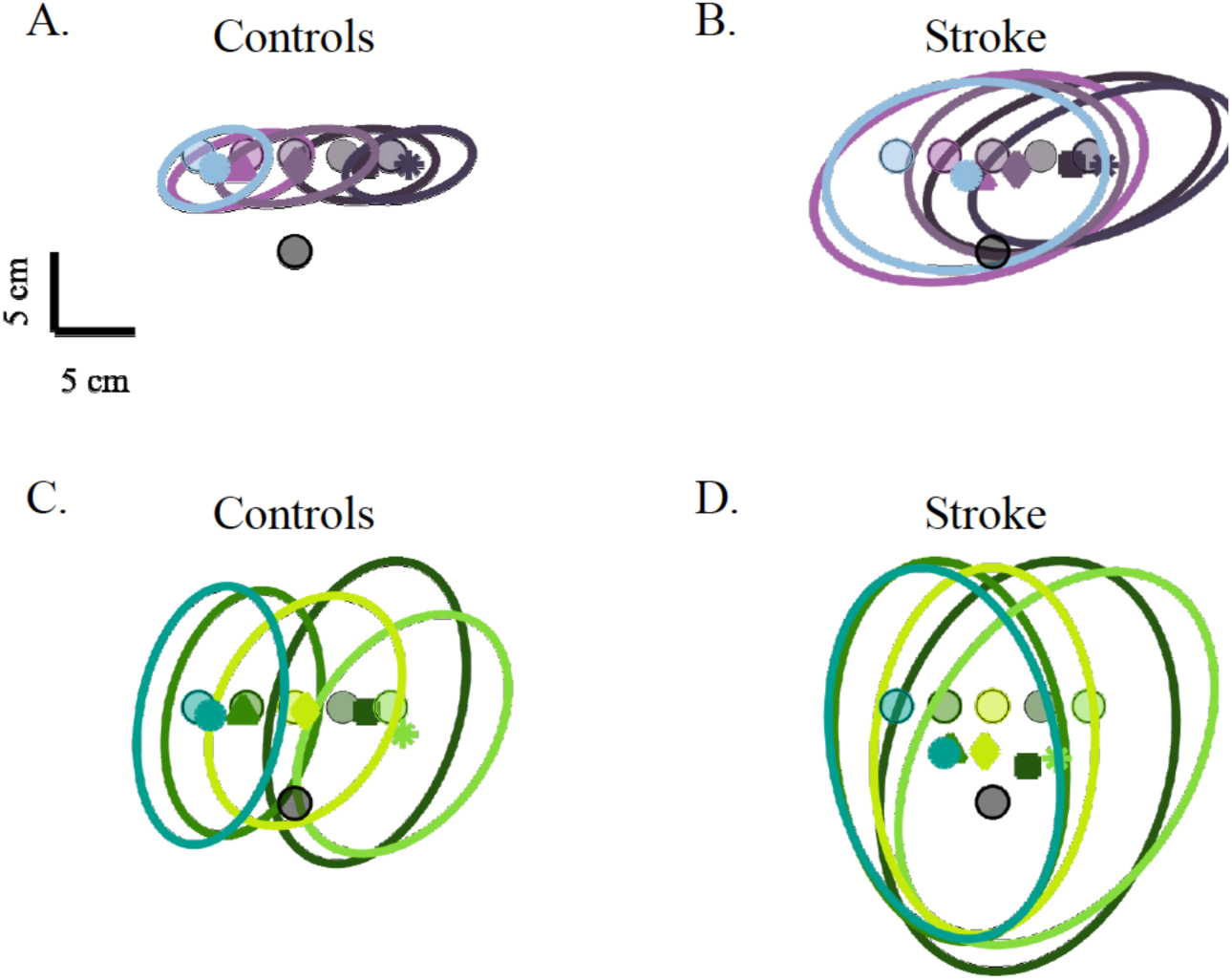
Average group level performance for arm matching (A, B) and eye matching (C, D) blocks from control (A, C) and stroke participants (B, D). Circular markers indicate the five target locations, with shape markers color coordinated to each target indicating the average Hand-EPE (A, B) and Eye-EPE (C, D). Each ellipse represents one standard deviation of the group data.

To determine if there were group level differences, we compared Hand-EPE and Eye-EPE between stroke participants and controls. Stroke participants showed significantly larger Hand-EPE compared to controls during arm-matching blocks (Figure 4A, Controls: 3.3 ± 1.4 cm; Stroke: 7.0 ± 4.1 cm; p < 0.001, CLES = 85.25). Similarly, we found that stroke participants had significantly larger Eye-EPE compared to controls, suggesting that proprioceptive deficits can be captured not only when using the arm as the end-effector, but also when using the eye as the end-effector (Figure 4B, Controls: 7.2 ± 2.8 cm; Stroke: 11.0 ± 4.0 cm; p < 0.001, CLES = 79.50).

**Figure 4:**
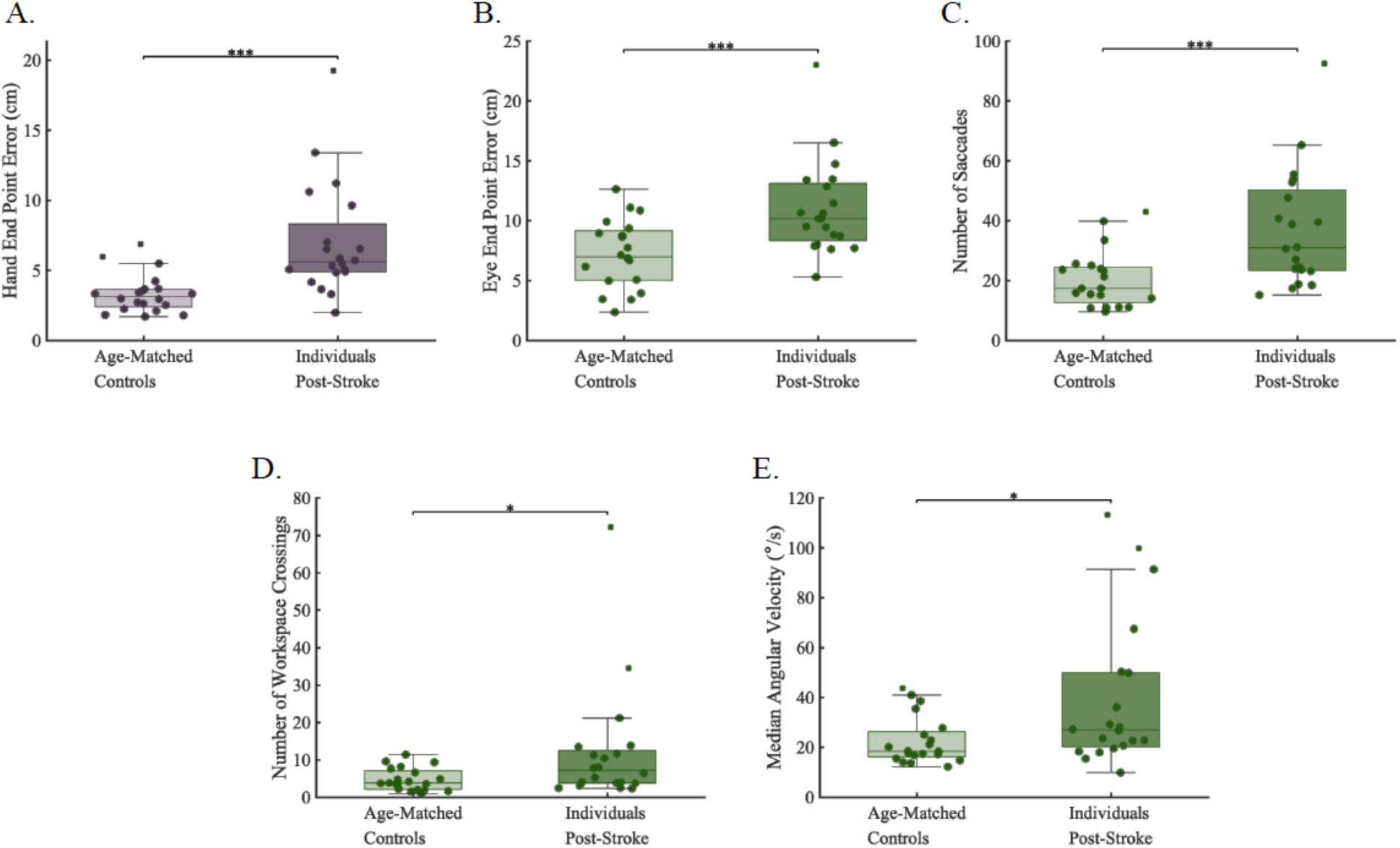
A). Individuals with stroke had significantly greater Hand-EPE during hand blocks compared to controls. B-E). Individuals with stroke had significantly greater Eye-EPE, Number of Saccades, Workspace Crossings, and Gaze Median Angular Velocity during eye matching blocks compared to controls. The eye data suggests that when using the more-affected limb as a reference, stroke survivors have significantly increased eye movement errors and increased attempts to use visual information to reference the more-affected limb. Box plots show median, upper and lower quartiles, and error bars indicate minimum and maximum values. *p ≤ 0.05, **p ≤ 0.01, ***p ≤ 0.001.

To better understand how impaired proprioception of the limb influences the execution of eye movements, we completed secondary analyses examining additional eye behavior kinematics. We found that stroke participants made a significantly greater number of saccadic eye movements over the course of a trial within the eye blocks compared to controls (Figure 4C, Controls: 20.4 ± 9.6; Stroke: 37.0 ± 19.6; p = 0.001, CLES = 78.25), a greater number of workspace crossings during the course of a trial (Figure 4D, Controls: 4.7 ± 3.0; Stroke: 12.1 ± 16.2; p = 0.011, CLES = 71.50). Lastly, we found that stroke participants had significantly faster eye movement velocities compared to controls (Figure 4E, Controls: 22.6 ± 9.7 °/s; Stroke: 40.0 ± 30.1 °/s; p = 0.017, CLES = 71.50). Finally, to better understand if errors in limb proprioception scale with error responses observed for eye movements (i.e., if individuals with large errors during limb matching also have large errors during eye matching), we examined the relationship between Hand-EPE and Eye-EPE. We found that there was a significant positive correlation between the amount of Hand-EPE and Eye-EPE for stroke participants (rho = 0.67, p = 0.001), but not for control participants (rho = 0.04, p = 0.45) (Figure 5), suggesting that poor proprioception of the limb can negatively impact estimated position of the eye.

**Figure 5:**
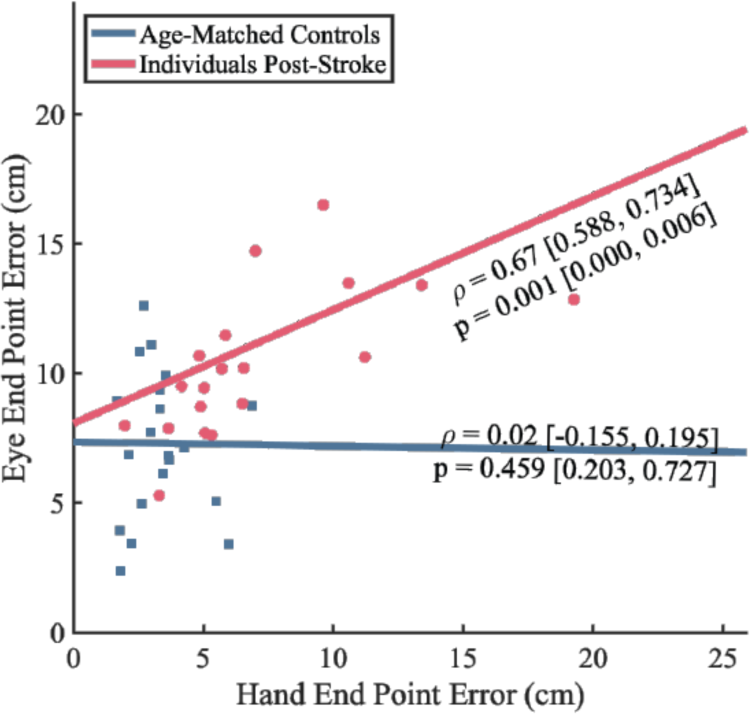
Spearman correlations for each group between Hand-EPE during the hand blocks and Eye-EPE during the eye blocks. Lines are the average of bootstrapped Ordinary Least Squares coefficients for each. Shown statistics are the median and 25th and 75th quartile of the bootstrapped distribution. The control group showed no correlation between Hand-EPE and Eye-EPE, while stroke participants showed a significantly strong positive correlation.

## Discussion

The goal of the current study was to determine if poor proprioceptive function of the limb carries over to perceptual estimates in the visual system for the execution of eye movements. Here, we observed that individuals with stroke had significantly impaired proprioception of the arm compared to controls, which is in agreement with previous work^5,6^. Newly, we observed that individuals with stroke also had impaired estimation of eye position when they were required to use the more-affected arm as a positional reference for localizing their eye movements. Further, we found that stroke participants had impairments in proprioceptive accuracy of the limb that was highly correlated to eye movement errors, an observation not observed in control participants. Overall, this work suggests that impairments in limb proprioception can impact and potentially interfere with the generation of eye movements when using proprioceptive information from the limb as a reference for guiding movement.

### Impairments in arm-based proprioception

The task used in this study is similar to previously established position sense paradigms that have been used to quantify upper limb proprioception in stroke survivors^5^. The primary difference between this task compared to previous methods is the horizontal orientation of targets. The purpose of the hand blocks in the study design was to compare proprioceptive accuracy of the limb against the eye blocks, which required participants to generate an eye movement to localize the position of the more-affected arm in the absence of visual information. A second purpose of the hand blocks was to replicate previous results to ensure that our task design was appropriately measuring upper limb position sense in stroke survivors. In agreement with this previous work, we found that the stroke participants included in our study had significantly increased Hand-EPE compared to controls. Additionally, this work detailed that ∼50% of stroke survivors experienced significantly impaired proprioception^5^. Here, we observed a slightly lower, but similar rate of impairment in those with chronic stroke, where we found that 8 of 20 (40%) stroke participants would lie outside the 95% normative range established by control participant values for Hand-EPE (Figure 4A).

### Impairments in eye-based estimates of limb location

To investigate how arm-based proprioceptive information can influence other sensory systems after stroke, we examined how movement of the eye could mirror perceived limb position in the absence of vision of the arm. Overall, we found that estimation of proprioception of the arm via the eye was highly disordered in stroke survivors. This included a significant increase in the magnitude of Eye-EPE when participants were asked to direct their gaze to the mirrored location of where they felt their arm, as well as considerable differences in our secondary metrics. Here, we observed that stroke survivors also made significantly more saccades, a greater number of eye movement crossings along the midline, and on average, had greater eye movement velocity compared to controls. These differences are likely attributable to impaired proprioceptive information of the more-affected arm that is being used as a reference generate oculomotor output. Recent work has suggested that the network responsible for the recovery and processing of proprioceptive information after stroke has broad reach beyond just primary somatosensory cortex^39–42^. Specifically, previous studies have noted that the posterior parietal cortex (PPC) plays an important role of integrating hand and visually-derived target information, and that damage to the posterior parietal cortex (PPC) can significantly disrupt this integration of information for action execution^43,44^. This suggests two possible explanations of these results, the first being that increases in eye-based proprioceptive error are due to impaired proprioceptive signaling due to damage to primary areas of proprioceptive processing, and/or damage that results in poor integration of proprioception with eye-based information^19,20,45–49^. To more fully understand the mechanisms underlying this phenomenon, future neuroimaging studies are necessary to confirm lesion-behavior relationships as well as functional contributions of these brain areas.

A surprising result that arose from the analysis of our secondary eye movement measures (angular velocity, number of workspace crossings, and number of saccades), was that all of these measures were elevated in the stroke group. During qualitative observation of data kinematics, some stroke participants clearly made repeated references with their eyes near the location of their stroke affected arm, despite having no visual feedback about the location of the arm. This manifested as an increase in the number of workspace crossings, which suggests that participants had difficulty making an accurate perceptual decision likely due to a noisy sensory reference which led to repeatedly referencing the general location of the both limbs, despite impaired sensation of the more-affected limb (Supplemental Figure 1). The negative impact of systemic noise on movement and perceptual decisions has been previously reported as leading to poor motor execution and can negatively impact transmission of sensory information to other sensorimotor areas of the brain^50,51^.

### Potential Limitations

To ensure that the increase in the magnitude of eye-movement-based errors observed in the stroke group was not the result of disordered saccadic control, we included visual fixation blocks that were interleaved with the hand and eye blocks during the experimental protocol. During these visual fixation blocks, we were able to capture naturalistic saccadic eye movements by simply having participants make a saccade to a visual target. Within the stroke group, we observed no relationship between level of proprioceptive impairment and the ability of stroke survivors to generate saccadic eye movements^34,52^ (Supplemental Methods and Results). However, we must note that a significant effect of age independent of degree of proprioceptive impairment was observed within the stroke group related to saccadic accuracy (Supplemental Figure 2). Another potential limitation of the current study is the potential influence of ipsilesional motor impairments on the ability of participants to perform the task. Previous work has shown that these impairments can occur in ∼40% of stroke survivors^53^. However, we have found that for examining bilateral proprioceptive matching tasks that removing these participants has negligible effects on reported results^6^.

## Conclusions/Implications

Overall, we found that proprioceptive impairments of the limb can disrupt the estimates required for accurate movement execution of the eyes. While the impact of proprioceptive impairments to overall levels of functional impairment after stroke are poorly understood, we find that sensory deficits of the limb may negatively impact eye-hand coordination when visual targets are not present. As this occurs for some, but not all stroke survivors that we tested, it is clear that to develop a more comprehensive understanding of the interactions between different modalities of sensorimotor function, it is likely necessary to adopt multi-method approaches to optimize treatment after stroke.

## Supporting information

Supplemental Methods and Results

## Acknowledgements

We wish to thank Kenna Gilley, Tami Wright, and Joanna Hoh for with subject recruitment, data collection, and conducting clinical assessments.

