## Supplemental Methods and Results for "Impairments in proprioceptively-referenced limb and eye movements in chronic stroke"

### *Data Analysis*

If eye-tracking data was missing from the entire trial, no cleaning or classification occurred (age-matched controls: 0 [0, 0] eye trials and stroke participants: 0.35 [0, 0] eye trials [values reported as mean and the $25^{th}$ and $75^{th}$ percentiles]).

### *Eye Movement Kinematics*

Before analysis occurred, eye movement data was cleaned, and saccadic thresholds were determined. To clean the eye movement data, we first removed blinks and gaze artifacts. Next, the data were low pass filtered at 20 Hz and then transformed from 2-dimensional-robotic-space to eye-based 3-dimensional spherical coordinates. Next, we used the derivatives of the spherical coordinates to compute yaw, pitch, and roll. To determine the saccadic thresholds, we used a previous method from our group[1]. This method first extract local velocity peaks from the gaze data. Then, a bimodal lognormal distribution is fit to the distribution of local velocity peaks using maximum likelihood estimation. Finally, an exponential function is used to determine the saccade velocity threshold (Equation 10a, from Singh et al.).

### *Visual Fixation Block*

To ensure that impoverish eye movements did not influence the ability of participants to properly execute the Eye Movement blocks, we interleaved Visual Fixation Blocks. The purpose of this was to ensure that throughout the task, participants had the ability to appropriately fixate on the given visual target that appeared. To quantify the quality of this fixation during these blocks, we quantified the percent of trials that participants foveated the visual target. Here, we found that stroke participants, on average, foveated significantly fewer visual targets compared to control participants (p = 0.001, CLES = 77.38). To determine whether this difference was directly due to stroke or another mitigating factor, we performed bootstrapped Spearman correlations on the percent of foveated trials as a function of age. Age has been previously found to negatively impact eye movement behavior, particularly related to response latencies and directional errors of saccades (Peltsch et al. 2011). Here, we found that within our group of stroke participants, age was the only reliable predictor of the percent of trials foveated (rho = -0.51, p = 0.02). No significant effects were found for level of proprioceptive impairment, general motor status (Fugl-Meyer), or cognition (MoCA). Further, we confirmed that our primary outcome measure for quality of eye movements (Eye-EPE) was not significantly correlated, nor a strong predictor of the percent of trials foveated (rho = 0.10, p = 0.46).

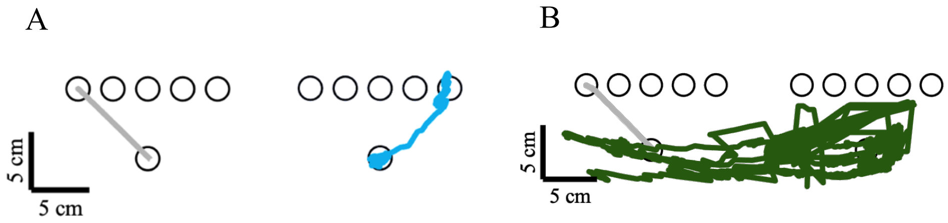


**Supplemental Figure 1.** Eye movement trajectories for a single trial during an eye movement block for a control participant (A) and stroke participant (B). A) During the course of a single trial, the control participant had zero workspace (midline) crossings as indicated from the blue trajectory from the start target position (bottom black outlined circle) to the upper right hand target. B) In contrast, the stroke participant had a high number of eye movement workspace crossings during a single trial performance, as indicated by the green line. Here, we observe that the eye movements of the stroke participant traverses the midline of the workspace numerous times over the course of a single trial.

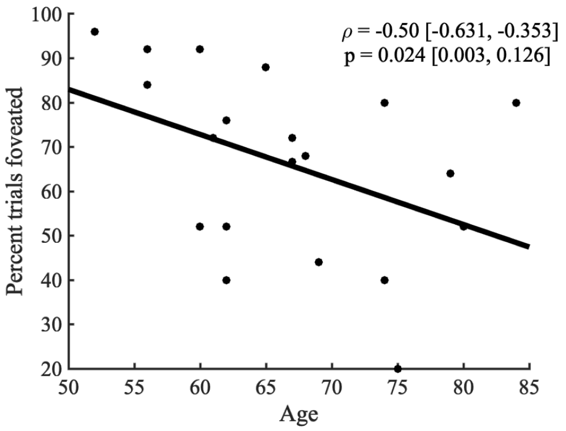


**Supplemental Figure 2.** To determine whether eye movements were impaired in stroke participants, we quantified the number of visual fixation trials that were foveated for each participant. Quantifying target foveation allows for determination of whether or not participants were appropriately fixating their eye movements on the targets that appears during the visual fixation blocks. We found that there was not a significant relationship between group and percent of trials foveate. This allowed us to determine that stroke was not a significant factor in the ability to foveate the visual targets, suggesting that eye movements were not impaired in our subject group as a result of stroke. However, upon examining the data related to the percent of trials foveated, we found that increased age was significantly related to a decrease in percentage of foveated targets during the visual fixation trials. The data in the figure above indicates the average percent foveation for each participant with stroke as a function of age. Data was fit via bootstrapped ordinary least squares. The above results are not surprising as effects of aging on eye movements has been previously established (Peltsch et al. 2011).
